# Utilizing combined spatial transcriptomics to elucidate localized immune responses within human coronary arteries throughout the progression of atherosclerosis

**DOI:** 10.1101/2025.02.20.639291

**Authors:** Joana Campos, Jack L McMurray, Michelangelo Certo, Ketaki Hardikar, Chris Morse, Clare Corfield, Melanie Weigand, Desley Neil, Pasquale Maffia, Claudio Mauro

## Abstract

Atherosclerosis is a complex inflammatory disease characterized by the accumulation of lipids and immune cells in the arterial wall, leading to the narrowing and stiffening of blood vessels. The involvement of both innate and adaptive immunity in the pathogenesis of human atherosclerosis is increasingly recognised. However, the spatial organization and specific roles of immune cells during the various stages of disease progression remain poorly understood, underscoring the necessity for additional research to elucidate their functions throughout the disease course. A better understanding of the immune response’s contribution to atherosclerosis progression could unveil novel therapeutic targets to mitigate plaque development and rupture, ultimately reducing the burden of cardiovascular events.

In this study, we utilised NanoString GeoMx^®^ and CosMx™ technologies to analyse serial sections of human coronary arteries from patients with varying degrees of atherosclerotic lesion severity. Our work consists of a series of investigations, and integrated findings from both the GeoMx^®^ and CosMx™ datasets, including pathway analyses, cell typing, and neighbourhood analysis. This workflow underscores the power of combining these spatial transcriptomics platforms to elucidate biological processes at the single-cell level, hence unbiasedly providing molecules and pathways of relevance to aid in the understanding of disease pathogenesis and assessing the opportunity of novel therapies.

## Main

Significant mechanistic advances in cardiovascular immunology have largely been driven by technological developments. It all began with the development of monoclonal antibodies, which enabled the use of immunofluorescence to map key immune cells in human atherosclerotic plaques for the first time in the 1980s^1–3^. Progress continued with the first flow cytometry analysis of the mouse aorta^4^, advanced imaging of arterial lymphocyte recruitment^5^, and the development of omics techniques such as cytometry by time of flight (CyTOF)^6^ and single-cell RNA sequencing (scRNA-seq)^7,8^. We now understand that the immune response in atherosclerosis is complex, with various immune cells and phenotypes detectable at different stages of the disease and in different layers of the arteries.

Recently, spatial transcriptomics has been applied to assign mRNA readouts to specific locations within carotid endarterectomy sections and coronary unstable lesions^9–12^. However, this powerful technique has not yet been used to achieve single-cell transcriptomic analysis of vascular tissue. Moreover, these samples provide limited insight into the early stages of disease progression and lack a comprehensive view of the perivascular immune mechanisms. To fully understand the immune response’s significant role in the plaque, adventitia, and perivascular space^13^, a comprehensive approach that includes multiple tissue layers is essential. This also involves longitudinal pathology studies, which underpin our approach to imaging coronary arteries at different stages of atherosclerosis development. Additionally, it is crucial to test various platforms, compare them directly, and integrate datasets to provide a holistic understanding.

In this study, we employed NanoString GeoMx^®^ Digital Spatial Profiler (DSP) and CosMx™ Spatial Molecular Imager (SMI) analyses on intact human coronary arteries at various stages of disease progression to probe the immunological processes underpinning atherosclerosis. These cross-sections of human coronary arteries, affected by atherosclerosis and obtained from explanted hearts of individuals who received heart transplants, include the full thickness of the vessel wall and surrounding adipose tissue with vasa vasorum.

Spatial transcriptomics has rapidly gained significance in many biomedical research fields, as it allows for transcriptomic analysis in a spatial cellular context. Our study is the first to report transcriptomic analysis of sequential human coronary artery sections using two combined spatial transcriptomic platforms. Using the GeoMx^®^ DSP platform, we conducted whole transcriptome analysis (∼18,000 transcripts) across regions of interest (ROIs) in coronary artery sections from three subjects with atherosclerotic lesions. ROIs covered infiltrating immune cells in the plaque and adventitia, as well as smooth muscle tissue. With the CosMx™ SMI platform, we accomplished single-cell transcriptomic analysis of the same tissues, using serial sections to those used for GeoMx^®^ DSP, where cells were profiled with a universal cell characterization panel consisting of 970 RNA probes (including 20 custom probes selected based on GeoMx dataset analyses).

Here, we describe a complementary workflow for transcriptomic analysis utilising these two platforms and demonstrate how combining these analyses provides a novel and powerful tool, which we anticipate will shape experimental approaches in translational and clinical medicine beyond atherosclerosis and cardiovascular disease.

### A spatial profiler approach in human atherosclerotic lesions

For this study, we accessed a cohort of human coronary artery samples from subjects who underwent heart transplants. These samples contained the full circumference and thickness/all layers of the artery, along with adjacent tissues. The samples show atherosclerosis progression from near-normal vessels (with adaptive intimal thickening but no atherosclerosis, ≤AHA type 3) to early intermediate (eccentric intimal thickening with early atheroma showing foam cells/extracellular lipid), late intermediate (increased size of the atheromatous plaque with luminal encroachment, development of a cap, and formation of a plaque shoulder), and severe lesions (large atheromatous plaques with marked narrowing of the lumen and increased fibrosis within the plaque, >AHA type 3), (Fig.1a). Sections from this cohort were stained with morphology markers (SYTO13, CD45 and CD4), to aid ROI placement, and hybridized with ∼18,000 barcoded RNA probes, which cover the whole human transcriptome (Fig. 1b).

**Figure 1.**
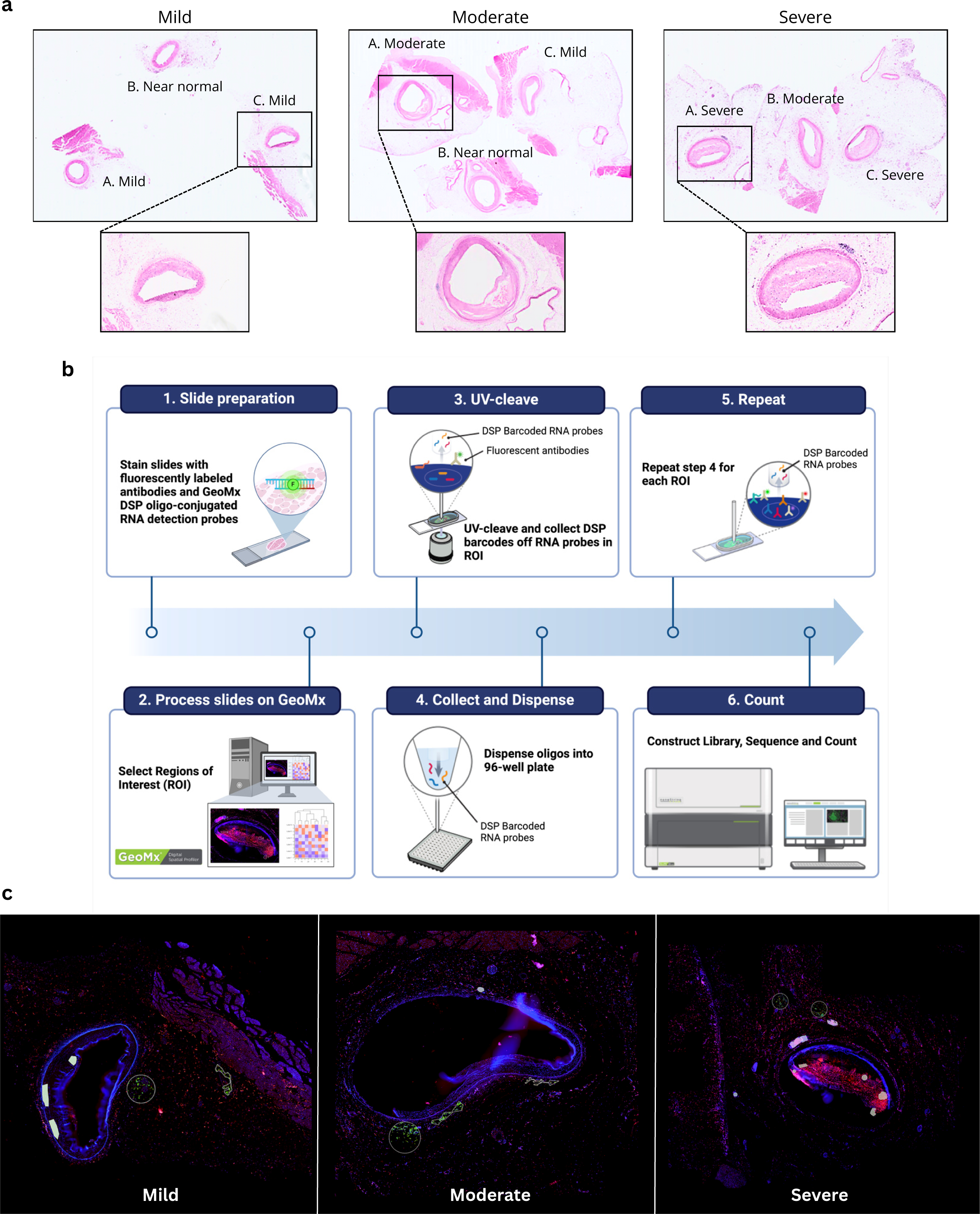
Histological Analysis and Workflow for GeoMx. **a,** Representative haematoxylin and eosin (H&E) stained sections showing arterial lesions of varying severity (Mild, Moderate, Severe). Specific regions are highlighted at a higher magnification to reveal morphological differences across lesion severity, indicating progressive changes from near-normal to severe. **b,** Workflow for spatial transcriptomics using GeoMx DSP: slide preparation with morphology markers and ∼18,000 oligo-conjugated RNA probes; selection of ROIs in each sample analysed; cleavage with UV light of barcodes from RNA probes; collection and release of collected barcodes onto a 96-well plate for all selected ROIs; generation of a cDNA library for Next Generation Sequencing and upload of resulting sequencing data onto the GeoMx DSP. **c,** Fluorescent imaging of arterial lesions with varying severities, reflecting those shown at higher magnification in A. Fluorescent imaging is a prerequisite for choosing Regions of Interest (ROIs) for downstream profiling by GeoMx. Samples were stained with SYTO13 (nuclear dye, blue), CD45 (pan-leukocyte marker, yellow) and CD4 (T cell subset marker, red). Representative ROIs chosen for downstream spatial profiling are indicated by white circles.

ROIs (n=55) were placed throughout the available coronary arteries to sample immune cells (CD45+CD4- and CD45+CD4+) infiltrating various layers of atherosclerosis-affected vessels, i.e., the plaque, adventitia, and smooth muscle cell layer (Fig. 1c). Approximately 3,000 genes passed QC and were further analysed.

### Infiltrating immune cell transcriptomic heterogeneity unveiled with the whole transcriptome atlas

As an initial approach, we performed unsupervised hierarchical clustering of immune cell segments across the plaque and adventitia (early [≤AHA type 3] and advanced [>AHA type 3] lesions), which revealed distinct transcriptomic profiles across the dataset (Fig. 2a). Comparing immune cells from different layers of the atherosclerotic vessel, we found most differentially expressed genes in the adventitia as compared to the plaque. In the adventitia, PLA2G2A, C7, and C3 were upregulated, while SULF1, FN1 and SPP1 were more prominent in the plaque (Fig. 2b).

**Figure 2.**
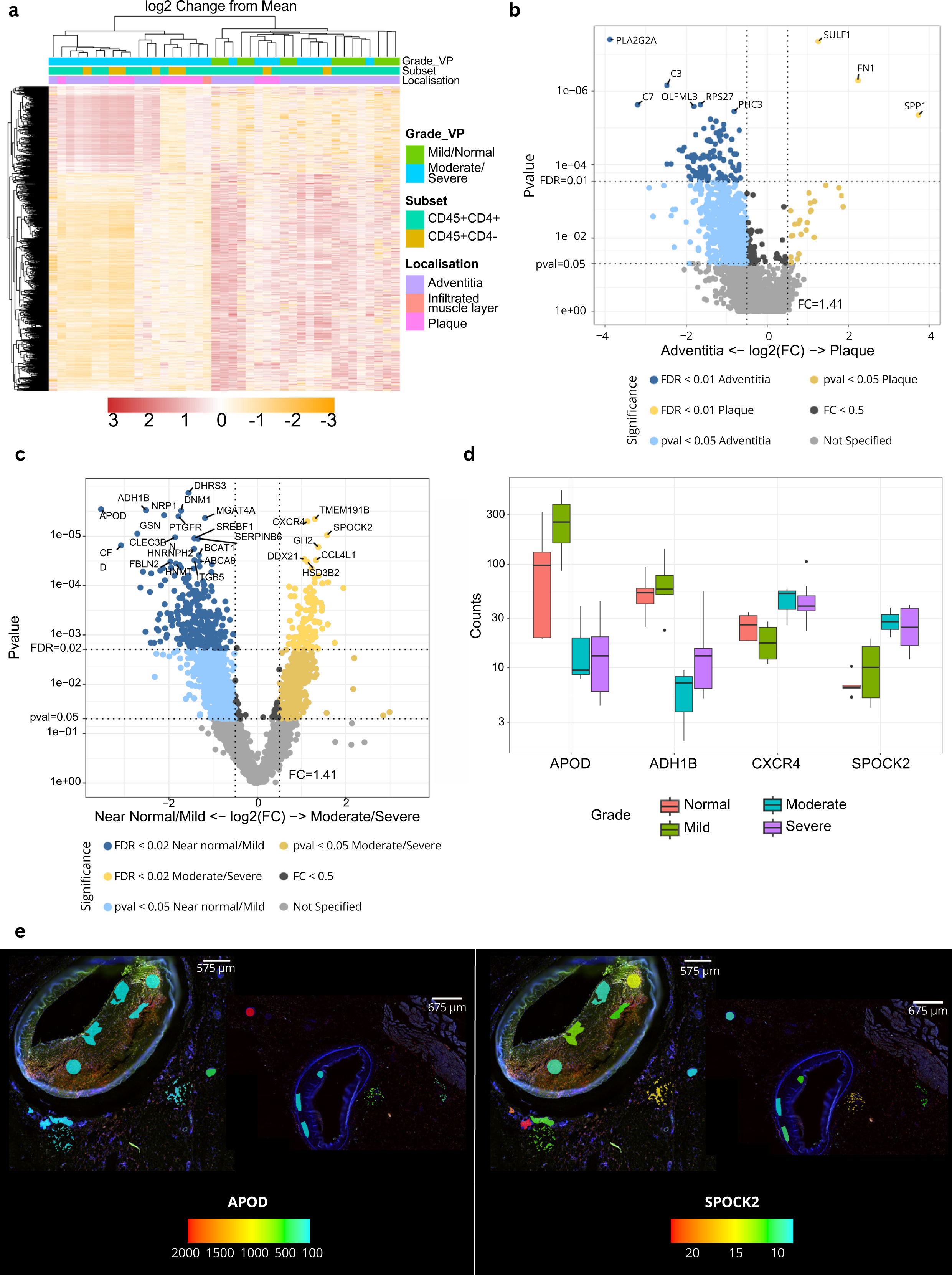
Spatial Transcriptomic Profiling of Arterial Lesions using GeoMx. **a,** Heatmap demonstrating the log2 fold change from the mean of gene expression of all targets post-QC. Heatmap is annotated by lesion severity (mild/normal and moderate/severe), CD45+ subset (CD45+CD4+ and CD45+CD4-), and tissue location (adventitia, infiltrated muscle layer and plaque). Scale bar represents log2 change from mean gene expression. **b,** Volcano plot exhibiting differentially expressed targets between CD45+ segments located in the adventitia and in the plaque. **c,** Volcano plot displaying differentially expressed targets between CD45+ segments in the adventitia of early lesions (near normal/mild) and those located in the adventitia of advanced lesions (moderate/severe). **d,** Box plots displaying counts of the top two differentially expressed genes in early (APOD, ADH1B) vs advanced lesions (CXCR4 and SPOCK2), as shown in C. **e,** Overlays of APOD and SPOCK2 onto the immunofluorescence scan of a severe (left) and a mild (right) lesion highlighting the targets’ expression in each profiled ROI. Overlay generated using the SpatialOmicsOverlay R package. Scale bar represents gene normalised counts.

To further investigate these differences, we grouped the lesions into early (near normal and mild) and advanced (moderate and severe). Immune cells in the adventitia of early vs. advanced lesions displayed diversity in gene expression, with several genes significantly enriched at different stages of disease progression (Fig. 2c). APOD and ADH1B were upregulated in early lesions, and CXCR4 and SPOCK2 in advanced lesions (Fig. 2d). Using a publicly available R package provided by NanoString (SpatialOmicsOverlay version 1.4.0), we were able to transpose gene expression levels, as exemplified by APOD and SPOCK2, onto immunofluorescence scans of mild and severe lesions, showing increased gene expression in the adventitia of advanced lesions (Fig. 2e).

### Whole transcriptome atlas informs pathway analysis

The GeoMx whole transcriptome atlas provides one of the most comprehensive tools to performing pathway analysis. To investigate the enrichment of differentially expressed genes into biologically meaningful pathways, we performed gene set enrichment analysis (GSEA) using the Biological Processes section of the Gene Ontology (GO) database. Interestingly, significantly enriched pathways were identified in the adventitia of near normal/mild vessels compared to moderate/severe vessels (Fig. 3a, Supplementary Table 1), suggesting a de-differentiated/stem-like immune cell phenotype in severe lesions. Pathways related to Actin Filament Organisation and Sterol Transport were among the most significantly altered (Fig. 3b). However, whilst the Actin Filament Organisation pathway is overall significantly increased in early lesions, a subset of targets assigned to this pathway is upregulated in advanced lesions, highlighting the possibility of further investigations from these pathway analysis outputs (Fig. 3c).

**Figure 3.**
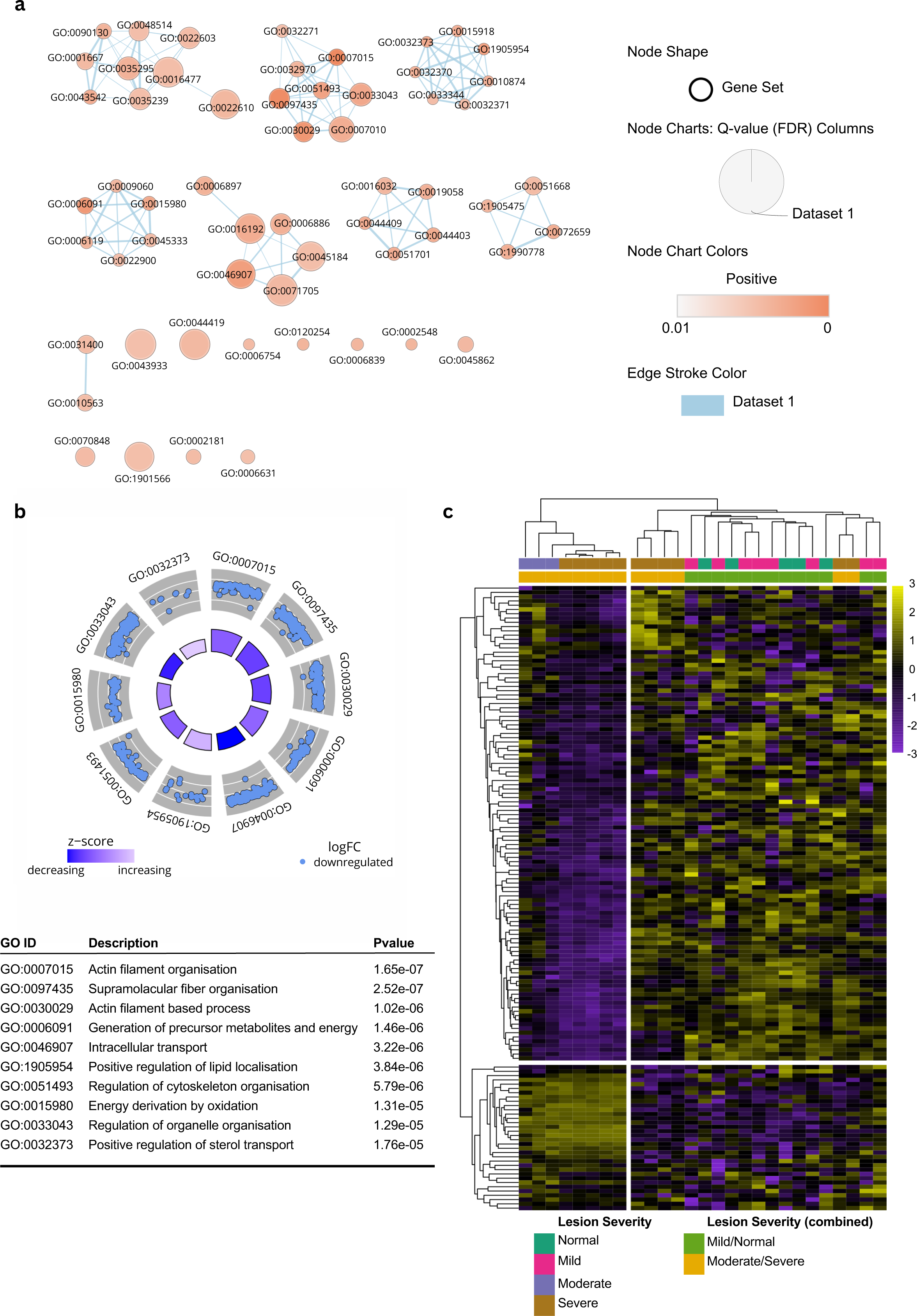
Pathway Analysis from Spatially Resolved GeoMx Dataset. **a,** Gene set analysis using the Biological Processes section of the Gene Ontology database reveals 58 differentially enriched pathways in CD45+ segments in the adventitia of early compared to advanced lesions. The enrichment map was produced with Cytoscape where each the size of each circular node represents the size of an enriched pathway. Jaccard similarity scores evaluated gene overlap between pathways whereby a thicker line represents increasing similarity. Finally, nodes were colours by the degree of enrichment (FDR value), whereby a deeper red indicates a more significant pathway. Only genesets with an FDR < 0.01 are shown. **b,** GOCircle plot representing the top 10 differentially enriched pathways. Scale bar represents z-score calculated as (up-regulated genes – down-regulated genes)/√total genes, indicating over (high z-score) or underrepresentation (low z-score) of genes in a pathway. Outer circle of the plot displaying a scatter plot for each pathway of the logFC of the assigned genes (each gene represented as a dot in the scatter plot). Inner circle plotting a bar plot where bar colour corresponds to z-score (blue, decreased; white, increased; and lavender, unchanged) and bar size reflects degree of significance (log10-adjusted P-value). Description and p values for top 10 enriched pathways also displayed. GOCircle plot generated using GOplot 1.0.2 R package. **c,** Heatmap displaying log2 change from mean of targets included in the top enriched pathway (GO:0007015 Actin Filament Organisation).

### Single cell spatial transcriptomic resolution using CosMx Spatial Molecular Imager

We extended our spatial transcriptomics workflow to analyse serial sections from the same samples used in the GeoMx^®^ study, using NanoString’s latest spatial profiler, CosMx. This is a single-cell spatial imager with a similar slide preparation method to that required for GeoMx^®^ studies (Fig. 4a). Oligo-conjugated RNA probes are hybridized to tissue sections on glass slides, which are then imaged within the instrument following a series of reporter cycling steps. For this study, coronary artery sections were profiled with the CosMx^™^ Universal Cell Characterization RNA panel, which contained a core of 950 probes and an additional set of 20 custom probes (specific to this study and informed by the GeoMx^®^ DSP study). Fields Of View (FOVs, n=235) were placed to select areas of interest to be imaged and profiled. In this study, FOVs were placed in such way that the full thickness of the coronary arteries, including the plaque (when present), the media, the adventitia and infiltrating immune cells, were analysed (Fig. 4b). This represents a significant advantage by CosMx^™^ over GeoMx^®^’s limited ROI area.

**Figure 4.**
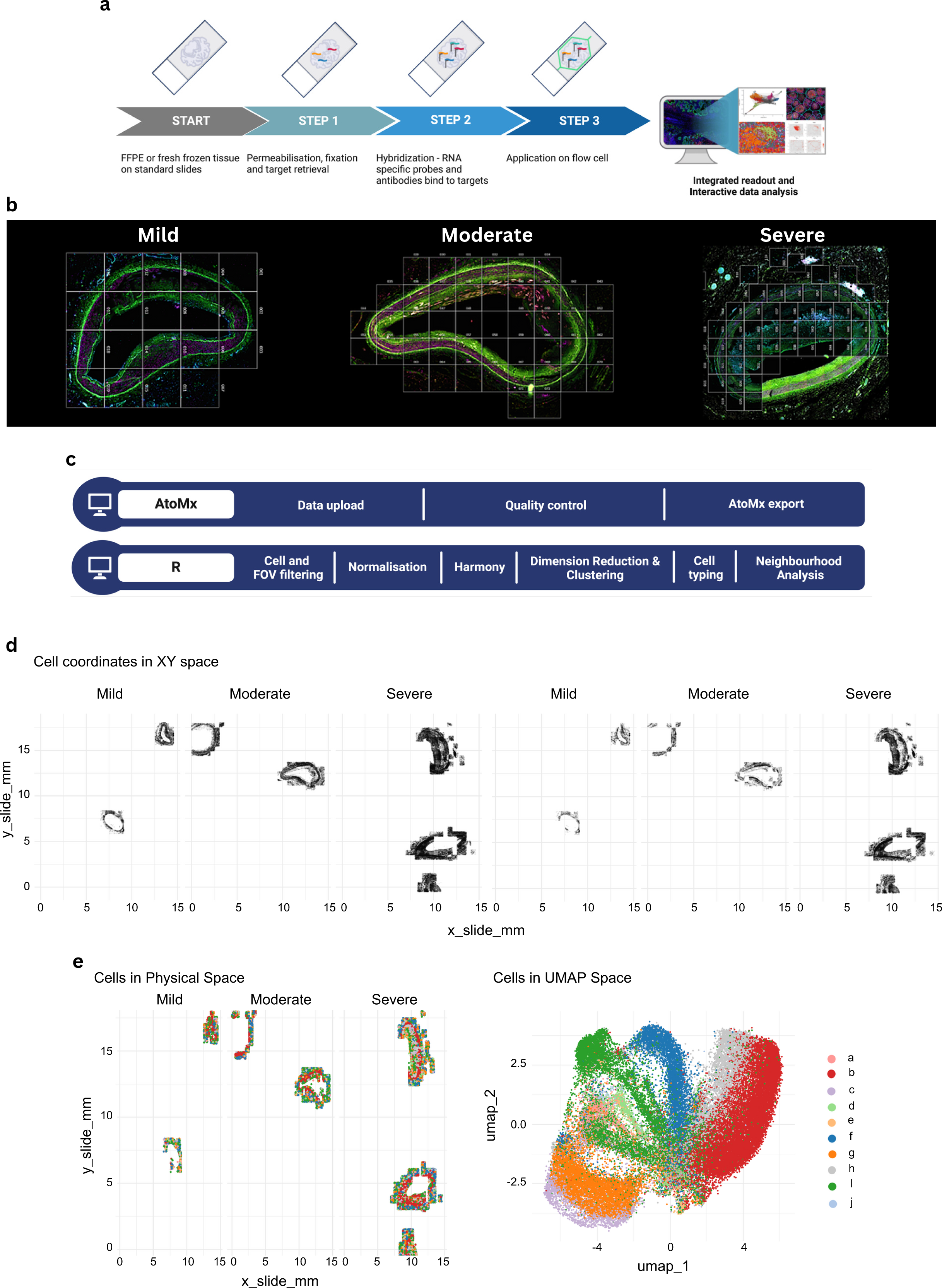
Workflow for single cell spatial transcriptomics using CosMx SMI platform. **a,** Schematic representation of steps involved in CosMx studies. **b,** Immunofluorescence scans generated by CosMx of representative mild, moderate and severe lesions, depicting FOV placement. Sections were stained with DAPI (nuclear staining, not visible), B2M/CD298 (cell membrane markers, cyan), PanCK (pan-epithelial cell marker, green) and CD45 (pan-leukocyte marker, red). **c,** Data analysis for CosMx was split between AtoMx and R; diagram highlights which parts of the process were performed on which platform. **d,** Plots displaying cells in the slide physical space for each lesion severity analysed. Left panel: all cells included in the dataset before cell and FOV filtering step. Right panel: cells remaining in the dataset post cell and FOV filtering. **e,** Complete post-filtering dataset shown as cells in physical space (left panel) and as a UMAP plot following unsupervised cell typing.

The current workflow for CosMx^™^ studies includes an automatic data push to AtoMx Spatial Informatics Platform (SIP), a NanoString cloud platform for dedicated data analysis of CosMx^™^-generated datasets. However, as of the writing of this article, AtoMx SIP remains a work in progress and does not yet provide a comprehensive solution for analysing complex datasets like those produced by CosMx^™^. Therefore, our use of AtoMx SIP in this study was limited to data upload from the CosMx^™^ instrument, quality control (QC), and data export to an AWS bucket. For downstream analyses, we employed publicly available R packages, resulting in a bespoke pipeline for analysing CosMx^™^ datasets (Fig. 4c).

One key addition to our CosMx data analysis pipeline is a cell and FOV filtering step, which excludes any cells or entire FOVs that do not meet the specified QC metrics. We believe that the data trimming resulting from this filtering step (Fig. 4d) is essential for any downstream analyses and should not be considered optional, as only data that successfully passes QC is carried forward. Overall, 970 genes passed QC and were further analysed.

We initially performed unsupervised clustering of the complete post-filtered dataset, which resulted in 10 distinct cluster assignments (Fig. 4e). However, these clusters were not assigned to specific cell types or subsets due to the unsupervised nature of the approach.

### Single cell typing provided by Census

We used a Census-automated cell annotation tool, which is pre-trained on Tabula Sapiens dataset for vasculature, to perform cell typing on the coronary arteries profiled in this study. This approach led to the identification of 11 distinct cell types: B cells; CD4+ memory T cells; CD4+ helper T cells; CD8+ cytotoxic T cells; two subsets of endothelial cells; fibroblasts; macrophages; myofibroblasts; and two subsets of smooth muscle cells (Fig. 5a). In the left panel of Figure 5a, all cells (represented as individual dots) are displayed in both their physical space within the tissue and the flow cell. In contrast, the right panel shows the cells in UMAP space, highlighting not only cell type enrichment but also transcriptomic similarity among different cell types.

**Figure 5.**
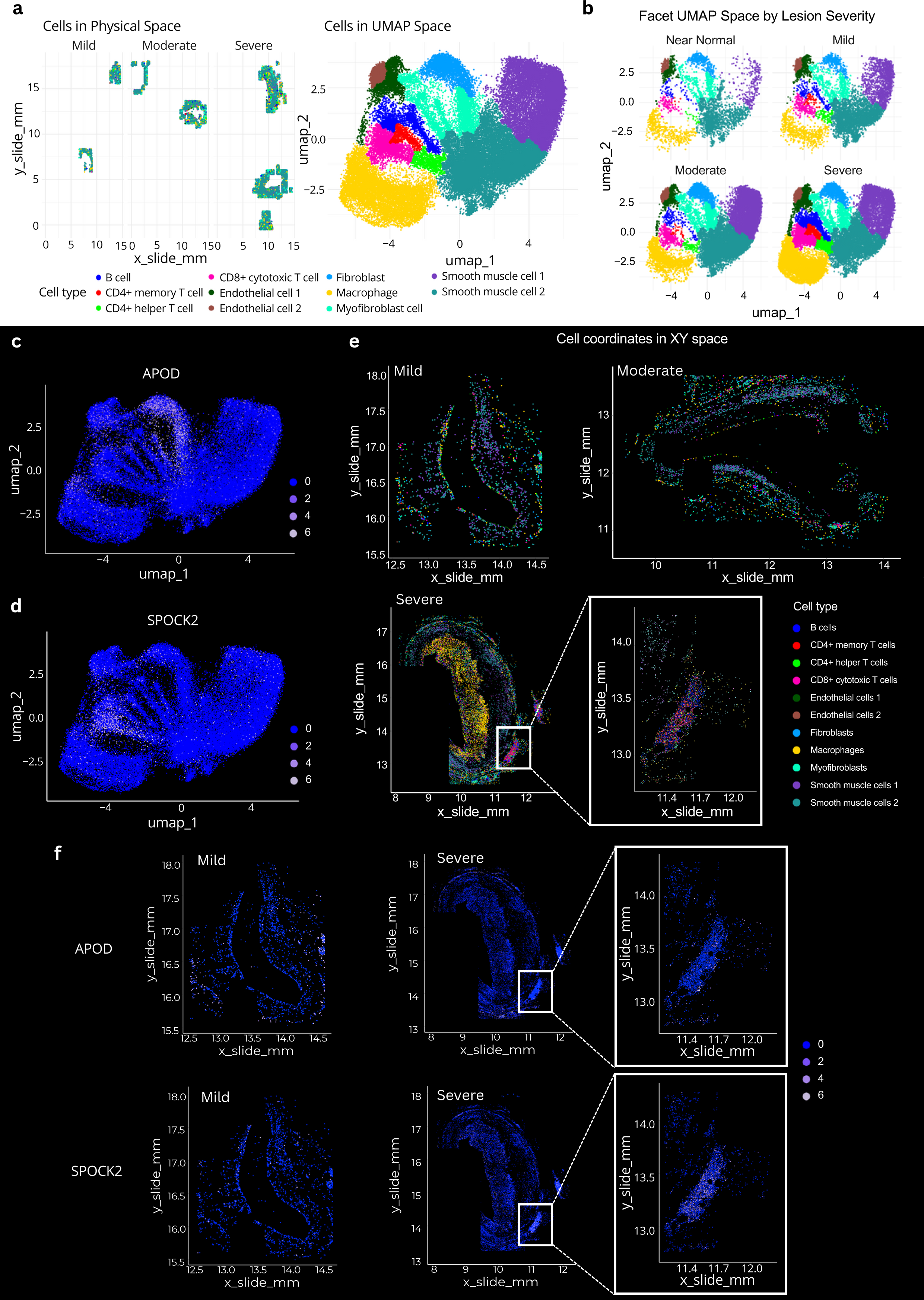
Single cell typing and target expression in CosMx datasets. **a,** Single cells from the entire post-QC dataset are identified in both the slide physical and UMAP spaces by their cell type, following supervised cell typing using Census vasculature signature. **b,** UMAP plots for the dataset split by lesion severity. **c,d,** APOD (C) and SPOCK2 (D) expression in individual cells represented in the entire dataset UMAP plots. **e,** Representative mild, moderate and severe lesions showing single-cell typing *in situ*. An ATLO region is highlighted at a higher magnification to reveal its cellular composition available through supervised cell typing. **f,** Single cell APOD and SPOCK2 expression levels in representative mild and severe lesions (including an ATLO).

To assess the success of the cell-typing method, we carefully examined the immunofluorescence scans generated by CosMx alongside the cell-typing readout. The vessel’s media layer was typed as being composed of two types of smooth muscle cells, while the plaque was mainly identified as macrophage and CD4+/CD8+ T cell infiltrates (Supplementary Fig. 1). A single layer of endothelial cells was also identified lining the vessel lumen, in agreement with current knowledge of these tissues (Supplementary Fig. 1).

An enrichment of several cell types linked with disease progression was observed when the UMAP dataset was split by lesion severity (Fig. 5b). Smooth muscle cells 1 and 2 were significantly enriched in severe lesions compared to early lesions, demonstrating the phenotypic modulation of vascular smooth muscle cells (VSMCs), which can adopt various phenotypes influencing atherosclerosis development and progression in both beneficial and detrimental ways^10,14^. Similarly, myofibroblasts showed a noticeable increase as the disease advanced, consistent with findings from scRNA-seq of human coronary arteries^15,16^. In contrast to much of the traditional literature on atherosclerosis, but in line with recent studies^17^, CD4+ T helper/memory, CD8+, and B cells were most enriched in severe lesions compared to early ones. Furthermore, a distinct subset of endothelial cells was identified in advanced lesions, marked by an elongation of the endothelial cells from earlier lesions, which became more pronounced in moderate and severe cases. This is consistent with the emergence of a transcriptionally distinct subset of endothelial cells limited to advanced lesions^18^ (Fig. 5b).

Furthermore, CosMx allows resolution of gene expression within specific cell types. Using example genes and pathways from the GeoMx^®^ dataset (Fig. 2), we show that APOD was expressed by fibroblasts, myofibroblasts, endothelial cells, and macrophages across all severities (Fig. 5c), while SPOCK2 was primarily enriched in CD8+ and CD4+ T cells in advanced lesions (Fig. 5d). We then performed cell typing of the whole tissues for each lesion severity (Fig. 5e). The cell typing output in the representative severe lesion highlights a macrophage-rich plaque occupying a significant part of the vessel lumen, and an organised immune cell aggregate, mostly consisting of B cells and CD4+ memory/helper T cells, resembling arterial tertiary lymphoid organs (ATLOs) found in the perivascular space during experimental atherosclerosis^13^. Zooming in the ATLO associated with the severe lesion revealed that B cells, CD4+ T cells, and high endothelial venule cells, typical of ectopic lymphoid structures, were most enriched (Fig. 5e).

Finally, we plotted the same genes as in Figures 2e, 5c and 5d (APOD and SPOCK2) onto the whole tissue. As expected, APOD was mainly expressed in the media and adventitia, while SPOCK2 was primarily associated with immune cells, as evidenced by its expression in the ATLO (Fig. 5f).

### Cell quantification and neighbourhood analysis in CosMx datasets

Next, we quantitatively explored the CosMx^™^ dataset in terms of cellular density, gene expression, and cellular neighbourhoods. We first quantified the immune cell density across lesion severity. Normalising cell counts to the number of cells per FOV, we observed an increased immune infiltrate of all tested populations in the “severe” category (Fig. 6a). To further probe the microenvironment within lesions, we identified 10 cellular neighbourhoods based on the proximity (within 20µm) of cell types using k-means clustering. We then examined the spatial relationships of these neighbourhoods in both physical and UMAP space (Fig. 6b). Generally, the neighbourhoods consisted of a single cell type, as highlighted by the UMAP mapping of neighbourhood’s annotations onto the cell type UMAP from Fig. 5a, suggesting that most cell types reside in neighbourhoods with cells of the same annotation. To probe this further, we calculated the frequency of each neighbourhood across each of the lesion severities (Fig. 6c). While all neighbourhoods were present in all severity levels (Near Normal, Mild, Moderate, Severe), their frequencies varied. Neighbourhood 1 and 7 were particularly enriched in Near Normal lesions, while Neighbourhoods 3, 5, 9, and 10 were enriched in advanced lesions. Finally, we plotted a heatmap of neighbourhood compositions (Fig. 6d), showing that myofibroblasts and endothelial cells were enriched in Near Normal lesions, while immune cell populations such as macrophages (Neighbourhoods 5, 3, and 10) and B cells (Neighbourhood 9) were more prominent in advanced lesions.

**Figure 6.**
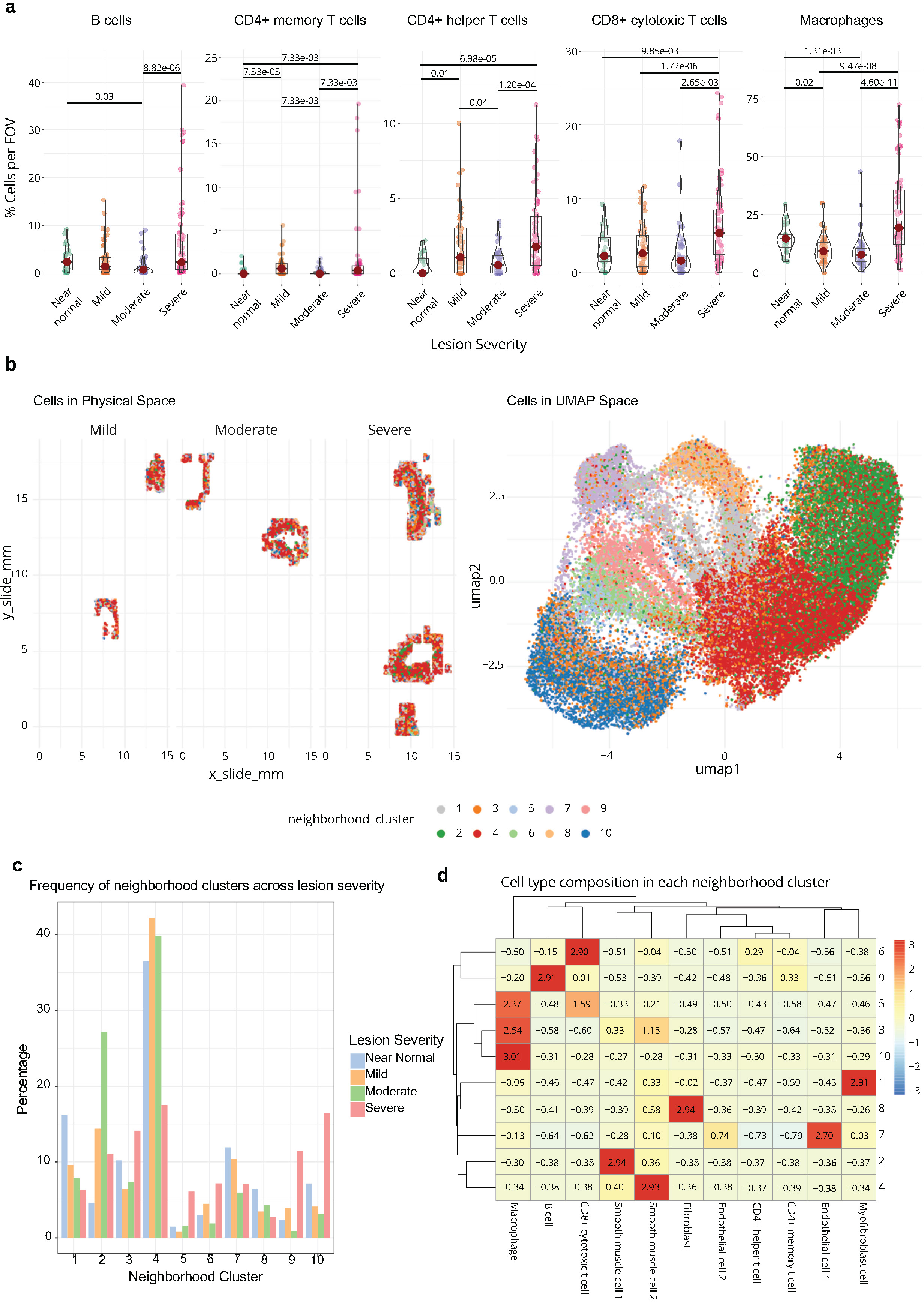
Quantifying cellular microenvironments across lesion severity. **a,** Violin plots depicting the percentage of different immune cell types (B cells, CD4+ memory T cells, CD4+ helper T cells, CD8+ cytotoxic T cells, and macrophages) per field of view (FOV) across lesion severity (Near Normal, Mild, Moderate, Severe). Only significant differences between cell densities across severities are shown (indicated by P values). **b,** Cellular neighbourhood clusters were defined using the frequency of annotated populations within 20μm of one another as input to clustering. Cellular neighbourhood clusters are shown in both physical space (left) and UMAP space (right). **c,** Bar plot showing the frequency of neighbourhood clusters across lesion severity (Near Normal, Mild, Moderate, Severe). Each bar represents a frequency of a cellular neighbourhood within a lesion severity. **d,** Heatmap of cell type composition within each neighbourhood cluster. The heatmap illustrates the relative enrichment or depletion of different cell types within each cluster, with colours representing z-score of enrichment (red – high, blue - low).

### Leveraging the integration of CosMx^™^ and GeoMx^®^ to explore targets beyond CosMx^™^ coverage

When it comes to target expression, GeoMx^®^ data is limited to a unique value per ROI/AOI within the tissue, as shown by the expression of ANXA2 mapped onto the immunofluorescence scan using SpatialOmicsOverlay (Fig. 7a). A significantly higher resolution can be achieved by CosMx^™^ when investigating the expression of the same target as per-cell specific expression levels are provided (Fig. 7a).

**Figure 7.**
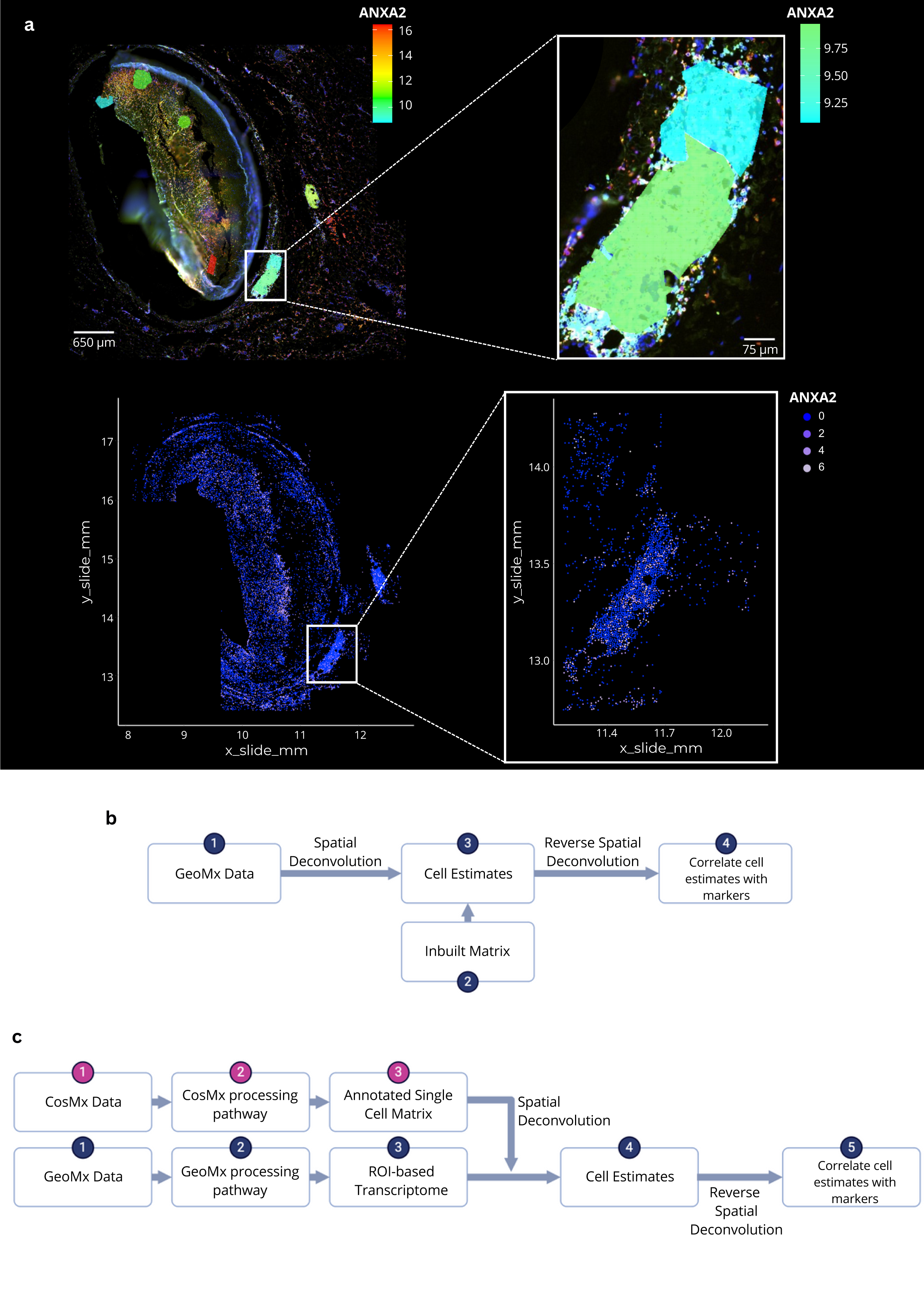
Integrating GeoMx^®^ and CosMx^™^ datasets: the current and the future. **a,** *In situ* expression of ANXA2 in a representative severe lesion in GeoMx^®^ (top panel) and CosMx^™^ (bottom panel). Higher magnification boxes show ANXA2 expression in an ATLO. **b,** NanoString-provided workflow for spatial deconvolution of GeoMx^®^ data. **c,** Proposed workflow for spatial deconvolution utilising a CosMx-generated single cell matrix (from the same tissue) to inform cell estimates.

Currently, NanoString provides algorithms for spatial deconvolution of cell type estimates from GeoMx^®^ regions. These algorithms rely on a previously generated and annotated cell type matrix to estimate cell type abundances within the ROIs (Fig. 7b). Whilst other experiments can assess the utility of these algorithms by nature of experimental design (ROIs selected for a specific cell type), we were presented with a unique opportunity in being able to directly compare all predicted cell types simultaneously by overlaying ROIs (GeoMx^®^) and FOVs (CosMx^™^) covering the same tissue areas. Furthermore, spatially deconvoluted GeoMx^®^ data can be improved by using an input matrix relevant to the tissue of interest, therefore, we hypothesized that this spatial deconvolution may be improved if we used a matrix derived from our CosMx^™^ dataset (Fig. 7c).

Firstly, we demonstrate a lack of correlation between the cellular estimates derived from the inbuilt matrix and the annotated ROI positions (Supplementary Fig. 2a). Here, regions chosen based on CD45+ CD4+ were largely being predicted to be populations consisting of Macrophages and NK cells, rather than CD4+ populations. To ameliorate this, we used our Census-annotated CosMx^™^ data to produce a reference matrix (Supplementary Fig. 2b) for spatial deconvolution of GeoMx^®^ ROIs. Whilst there seemed to demonstrate regions of gene specificity for the cell populations within the heatmap, after refining these genes to those retained post-filtering in the GeoMx dataset, these regions were all but lost – particularly in the CD4+ compartment (Supplementary Fig. 2c). The limited overlap of genes between our GeoMx^®^ and CosMx^™^ datasets (n = 366) likely hampered this attempt and future attempts will undoubtedly improve with the release of the 6,000plex CosMx panel. Therefore, unfortunately, spatial deconvolution using the CosMx^™^-derived reference matrix produced similar results to that of the inbuilt matrix (Supplementary Fig 2d).

Lastly, we hypothesised these estimates would allow us to perform reverse spatial deconvolution, whereby cell estimates would be used as input to estimate the gene expression of relevant targets within different cell populations. Using this approach, we expected to be able to compare the expression of any given target (present in the GeoMx^®^ whole transcriptome atlas panel but absent in CosMx^™^) across different lesion severities (Fig. 7c). However, unfortunately, due to the limited number of targets within the CosMx^™^ panel affecting the resolving of the GeoMx data, we could not confidently investigate the cell types expressing any given target.

This workgroup aimed at using CosMx*^™^* data to generate an annotated single cell matrix to be used as the source of single cell data to infer cell estimates in GeoMx^®^ studies (Fig. 7c). However, at this time with the resources available (i.e. panel plexity discrepancy between GeoMx and CosMx), this approach was not successful. Nonetheless, we believe that in the future the ability to generate a single cell matrix from serial sections of the same samples to those used in any given GeoMx study would be an invaluable and most accurate possible resource.

## Discussion

In this study, we employed a combined NanoString GeoMx^®^/CosMx™ approach to spatially resolve cell types and gene expression across different stages of atherosclerosis in human coronary arteries. To our knowledge, this is the first study to combine these two technologies for analysing coronary artery disease. Previously, similar spatial profiling approaches have been used exclusively in cancer research^19,20^.

Furthermore, the few studies that have presented spatial RNA sequencing data from cardiovascular tissues have focused mainly on human carotid endarterectomy specimens. These typically include only the luminal atherosclerotic plaque, with at most a thin strip of media, or unstable coronary artery lesions. Two of these studies utilized 10x Visium technology to uncover site-specific pathways driving human atherosclerotic plaque instability in endarterectomy samples^12^ and coronary arteries with an erosion phenotype^10^. In previous comparative analyses, the GeoMx^®^ platform was found superior for deep molecular profiling of known regions, making it more suitable for hypothesis-driven research compared to the 10x Visium platform^21^. More recently, the Resolve platform has been used with a limited panel of genes in carotid or femoral endarterectomy samples^11^.

Hence, our study is the first to apply a combined GeoMx^®^/CosMx™ approach to human vascular tissue, focusing on disease progression and immune response in the vascular environment. Additionally, because of the diverse clinical samples we utilized, it is the first to assess the full thickness of the arterial wall and surrounding adipose tissue in coronary arteries ranging from near normal to advanced atheromatous plaques. This allowed us to assess the immune cell phenotypes throughout disease progression in all specific layers of the arterial wall.

Our study yielded several significant findings, beginning with the identification of spatial contexts that influence the expression of key targets. We successfully performed both unsupervised and supervised cell typing across various disease stages, providing a comprehensive view of cellular and gene expression dynamics. Pathway analyses revealed that distinct cell types exhibit pathway enrichment based on their location, such as between the adventitia and plaque, as well as across different levels of disease severity. Additionally, unsupervised clustering of the datasets identified distinct transcriptomic signatures that change with disease severity and spatial context. UMAPs generated through CosMx^™^ revealed important differences in cell typing across severities, particularly the significant phenotypic variation in smooth muscle cells type 1 between early and advanced lesions. A new population of endothelial cells also emerged in advanced lesions, warranting further investigation. We have clearly demonstrated a significant expansion of adaptive immune cell subsets in advanced lesions. This finding is consistent with recent studies^17^ and reinforces the role of adaptive immunity in the pathology^22^. We also constructed cell-cell interaction networks and conducted neighborhood analyses, which identified unique cellular clusters based on cell type composition and spatial coordinates. Yet, deeper analyses of our combined GeoMx^®^/CosMx^™^ datasets in the near future will likely clarify the exact role of these immune cells in the pathogenesis of atherosclerosis and whether they can be targeted for therapeutic benefits.

Furthermore, we have attempted and believe we have a successful workflow for the integration of GeoMx^®^ and CosMx^™^ technologies once a whole transcriptome atlas panel is released for CosMx^™^, which would enhance our overall understanding of the cellular and molecular landscape. We discussed the potential to improve cell type delineation by expanding the panel of targets. Although single-cell sequencing often results in a significant loss of targets, increasing the panel plexity to the whole transcriptome on the CosMx^™^ platform will not render this workflow obsolete. This is because GeoMx sequencing provides deeper insights, allowing us to investigate rarer targets across different cell populations.

Recent advancements in integrating CosMx with single-cell RNA sequencing (scRNA-seq) have shown promising results^23^, offering valuable insights for understanding complex biological processes. By exploring how these integration methods enhance our comprehension of atherosclerosis, we can better position our findings within the broader context of current research and methodologies.

At the time of this study, the largest commercially available CosMx^™^ panel was the Universal Cell Characterization 1,000-plex. Currently, a 6,000-plex panel is available, and NanoString has publicly released data using the whole transcriptome panel for CosMx^™^, though it is expected to be commercially available by 2025. As a result, future GeoMx^®^ and CosMx^™^ integration will be feasible, providing the deepest cell typing and gene expression at the single cell level available, once the whole transcriptome panel becomes commercially available from NanoString.

In conclusion, our study provides a comprehensive framework for understanding cell type dynamics and immune responses in human atherosclerosis at a spatially single-cell resolution. The insights gained from this spatial transcriptomics approach offer valuable information for identifying and understanding novel pathways involved in the pathogenesis of human atherosclerosis, particularly in specific anatomical vascular niches and throughout its development and associated cardiovascular complications.

## Methods

### Samples

This study utilised 9 complete circumferential lengths of human coronary artery, including adventitia and surrounding adipose tissue, from 3 donors (1 female; 2 male), obtained from hearts removed at the time of cardiac transplantation. Three samples from the same formalin fixed coronary artery were embedded in each paraffin block and serial sections were cut from each block. The coronary artery cross sections varied within the same artery and the 3 donors were chosen to provide a range from near normal intima to severe atherosclerotic plaques. Ethics was obtained for method validation from the University of Birmingham Human Biomaterial Resource Centre (HBRC) RG_HBRCMV031.

### GeoMx^®^ DSP

For GeoMx^®^, 5µm FFPE sections were stained with a cocktail of morphology markers (SYTO13, CD45 and CD4) and hybridised with the Whole Human Transcriptome Atlas RNA panel (WTA, 18.677 probes). A total of 55 Regions of Interest (ROIs) were placed across the 9 cross-sections covering regions that were CD45+CD4^-^ or CD45+CD4^+^ in the adventitia or the atherosclerotic plaque. As controls, a few ROIs were drawn on CD45-CD4-regions (smooth muscle or nerve). The collected barcodes were processed and used to prepare a cDNA library for Next Generation Sequencing (NGS). The NGS readout data was converted from FASTQ to DCC format using the GeoMx NGS pipeline (version 2.0.0.16), and the data uploaded to the GeoMx^®^ DSP.

The whole slide scans were generated by the GeoMx^®^ DSP at the time of ROI placement and oligo collection and can be downloaded as OME. TIFF files were used for downstream analyses in packages such as the SpatialOmicsOverlay.

### GeoMx^®^ Data Analysis

Data Analysis was conducted on GeoMx^®^ Digital Spatial Profiler (DSP) version 3.1.0.194 following the User Manual recommended workflow and settings. Raw data from 55 ROIs profiled across 3 slides (each containing 3 artery sections as demonstrated in Fig. 1a) were subjected to technical and sequencing QC to assess the sequencing and overall quality of the data. Out of the 55 ROIs, 49 passed QC and were included in the downstream analysis. Of the 6 ROI segments that failed QC, 2 segments failed due to sequencing quality metrics, and the remainder due to low surface area and/or nuclei count. No segments were excluded at this step. Further, Probe QC was performed to identify negative control probe outliers, and none were reported. Filtering steps to exclude low expressing targets and segments were conducted which eliminated 14962 targets and 2 segments low in signal relative to the background. 3715 targets and 53 segments remained for further data analysis. These data were normalised using Q3 normalization and further subjected to unsupervised hierarchical clustering based on Pearson distance, Differential Gene Expression, and Pathway Analysis. Pathway analysis was performed in R (version 4.4.1) using the fgsea (fast gene set enrichment analysis) R package^24^ and the Biological Processes section of the Gene Ontology database. GO pathways were accessed through R using the msigdbr R package^25^. Pathways were filtered based on a minimum and maximum size of 15 and 500, respectively, and required to have a minimum of 20% of their targets covered by the GeoMx^®^ data. Pathways were visualised using GOCircle plots using the GOplot R package^26^, and Cytoscape^27^, using the R package RCy3^28^ and the Cytoscape plugin enrichment Map^29^.

### CosMx™ SMI

For CosMx™, serial 5µm FFPE sections (to those used for GeoMx) were hybridised with the CosMx Universal Cell Characterisation RNA Panel containing 950 core targets; this panel was then customised with 20 additional targets to cover genes and pathways highlighted by the GeoMx study. We placed 235 Fields of View (FOVs) across the 9 cross-sections to profile the full thickness of the artery layers, atherosclerotic plaques and immune cell infiltrates. RNA readout of placed FOVs included an incubation with the reporter pool followed by a reporter wash buffer to remove any unbound reporter probes. Imaging buffer was then released into the flow cell for imaging. Eight Z-stack images with a 0.8μm step size for each FOV were taken. Photocleavable linkers on the fluorophores of the reporter probes were released by UV illumination and washed off with strip wash buffer. The fluidics and imaging procedure was repeated for the 16 reporter pools, and the 16 rounds of reporter hybridisation-imaging were repeated multiple times to enhance RNA detection sensitivity.

### CosMx Data Analysis

Data analysis was initially performed using AtoMx Spatial Informatics Platform (SIP) version 1.2 following the User Manual instructions. However, initial segmentation was qualitatively deemed not suitable downstream processing, and cells were re-segmented with AtoMx SIP version 1.3. Post re-segmentation, cells were subjected to AtoMx’s standard quality control module default metrics, other than the average count for FOVs which was reduced to 35. 67% of cells passed these criteria, with 85% of FOVs being available for downstream processing.

Due to nuances with the AtoMx software, all further data exploration was performed outside of the AtoMx platform in the statistical programming language – R (version 4.4.1). A Seurat object was exported to Propath’s AWS S3 bucket and downloaded for further exploration. Data were first filtered for cells and FOVs passing quality control criteria and log normalised using the Seurat R package (version 5.1.0)^30^. Normalised data was scaled and subjected to linear (principal component analysis – PCA) and non-linear dimension reduction (uniform manifold approximation and projections – UMAP) for visualisation. Data were qualitatively checked for obvious batch effects and were harmonised to increase data integration for downstream clustering using Harmony^31^. A range of clustering mechanisms was evaluated, including Leiden (at a range of resolutions), InSituType^32^, and automatic cell typing using Census^33^. After evaluating each clustering mechanism by their tendency to fit the contours of the UMAP, Census was chosen as the clustering/annotation method for downstream analyses. Census is a pre-trained model using the Tabula Sapiens dataset, containing a “Vasculature” subset and allowed for automatic cell typing of the data. Downstream analyses revolved around the Lesion Severity variable, providing outputs such as cell density comparisons, neighbourhood analysis (using the RANN R package^34^), single gene enrichment analysis, and multi-marker enrichment analysis.

## Data availability

GeoMx and CosMx datasets are available under the GEO Accession database: GEOXXXXX

## Acknowledgements

We thank Propath UK for supporting this study. We thank HBRC at the University of Birmingham for providing the human tissues. We thank Nick Jones at the University of Swansea for critical reading of the manuscript.

## Authors’ Contributions

Conceptualization: J.C., J.L.M., M.C., D.N., P.M. and C.Mauro.

Methodology: J.C., J.L.M., M.C., K.H., M.W. and D.N.

Investigation: J.C., J.L.M., M.C., K.H., C.Morse, C.C., M.W. and D.N.

Formal Analysis: J.C., J.L.M., K.H.

Writing - Original Draft: J.C. and C.Mauro.

Writing - Review & Editing: J.C., J.L.M., M.C., M.W., D.N., P.M. and C.Mauro.

Supervision: J.C. and C.Mauro.

Funding Acquisition: C.Mauro.

All authors revised the manuscript critically and approved the final version.

## Ethics and Declarations

J.C., J.L.M., K.H., M.W., C.Morse and C.C. are employees of Propath UK. The remaining authors declare no conflicts in relation to this study.

---

**Supplementary Figure 1.**
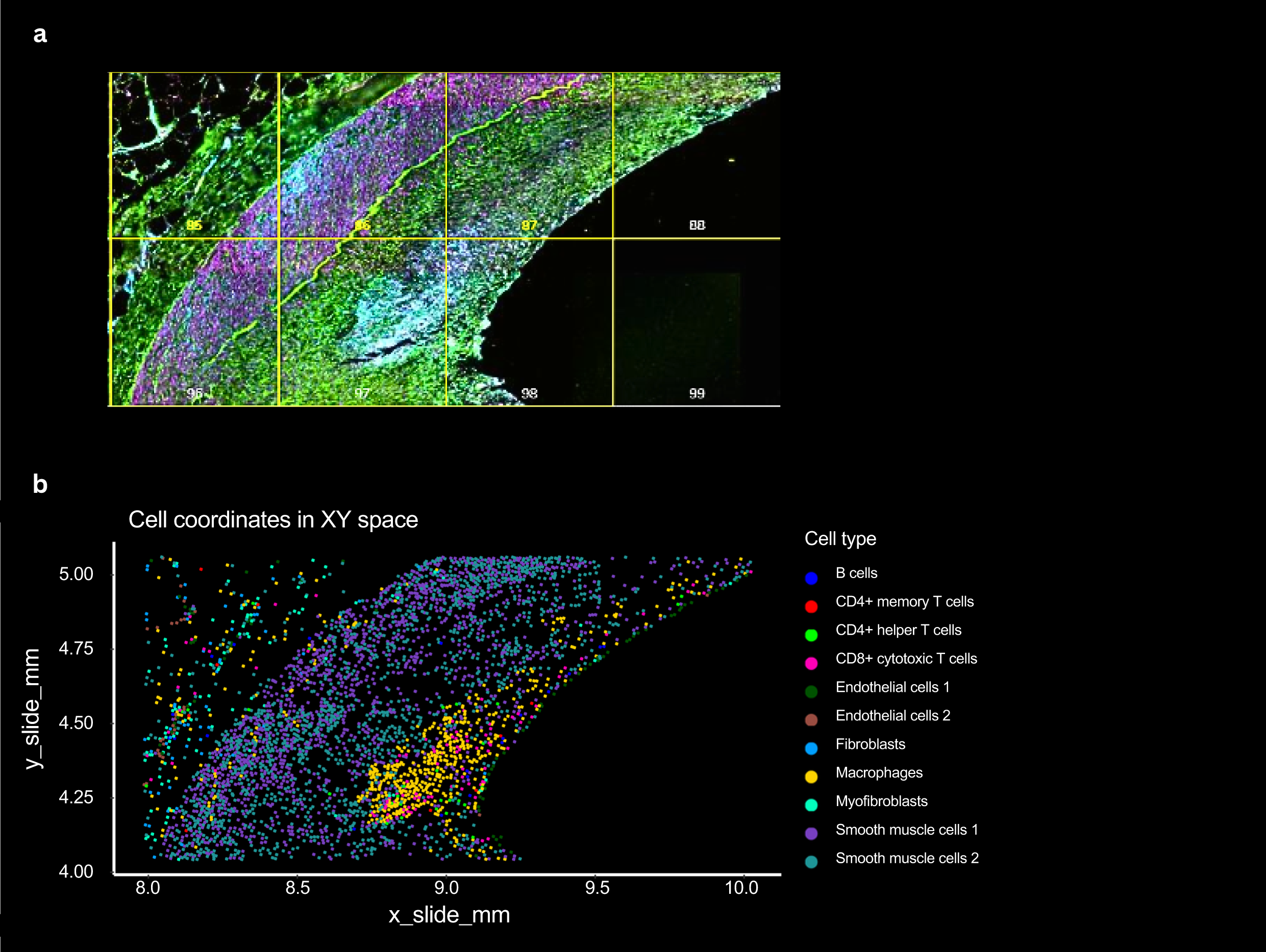
Immunofluorescence scan generated by CosMx™ showing the full thickness of the coronary artery **(a)** and respective cell typing equivalent following supervised cell typing using Census vasculature signature **(b)**.

**Supplementary Figure 2.**
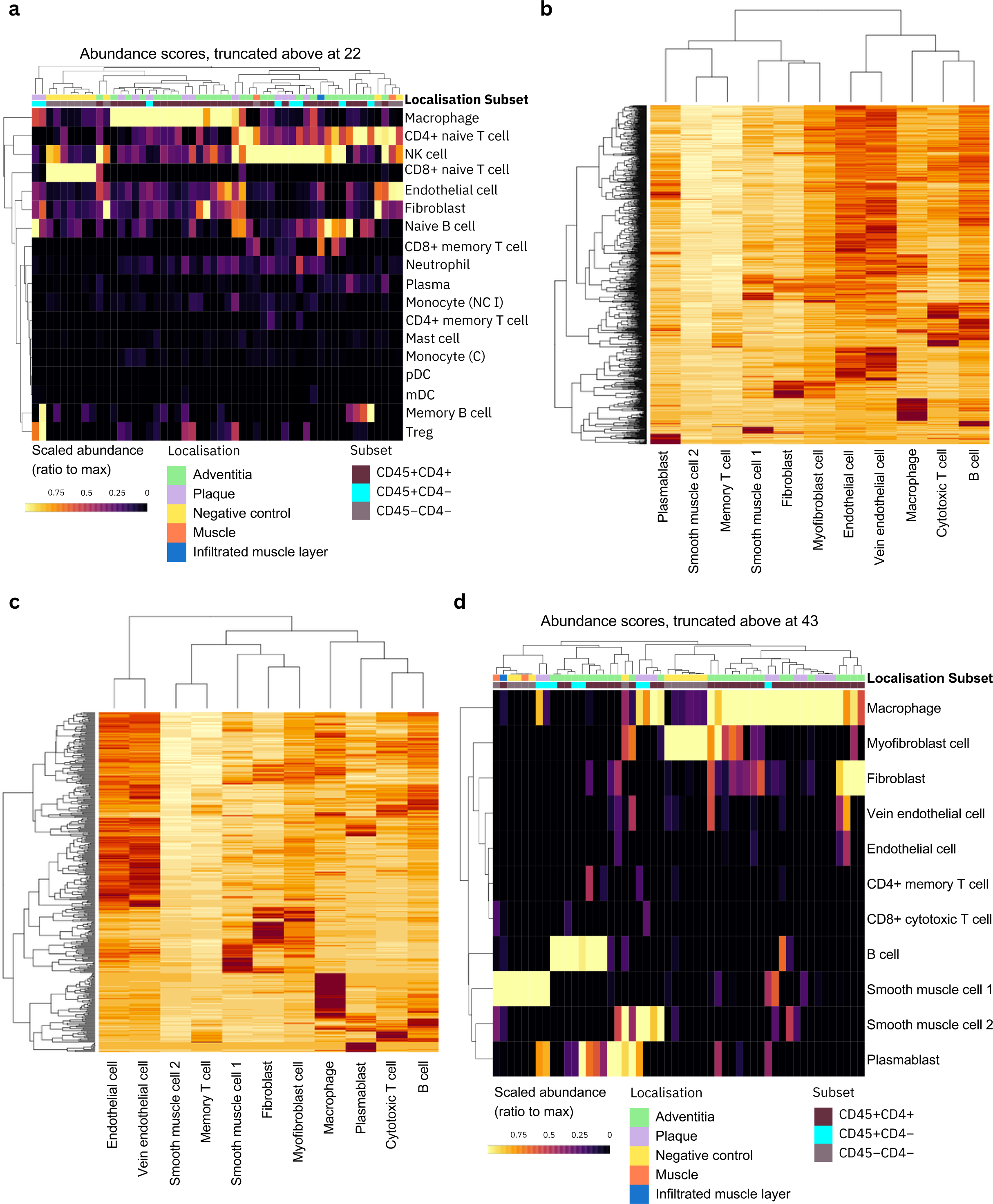
**a,** Heatmap of spatial deconvolution estimates using the inbuilt GeoMx reference matrix “safeTME”. Scaled abundances are shown as a ratio to the maximum value are displayed across five tissue localisations (adventitia, plaque, negative control, muscle, and infiltrated muscle layer). “Subsets” correspond to ROIs segmented on the GeoMx platform and are highlighted. Cell types present in the safeTME matrix are indicated as rows. **b,** Heatmap of the CosMx-derived cell signature matrix. Census-annotated cell populations from the CosMx are represented as columns with rows representing genes on the CosMx platform. Genes are scaled from red to white, with red indicating a higher expression. **c,** Heatmap of the genes present in the GeoMx dataset from CosMx-derived cell signature matrix. **d,** Heatmap of spatial deconvolution estimates using the CosMx-derived matrix. Cell types present in the CosMx-matrix are indicated as rows.

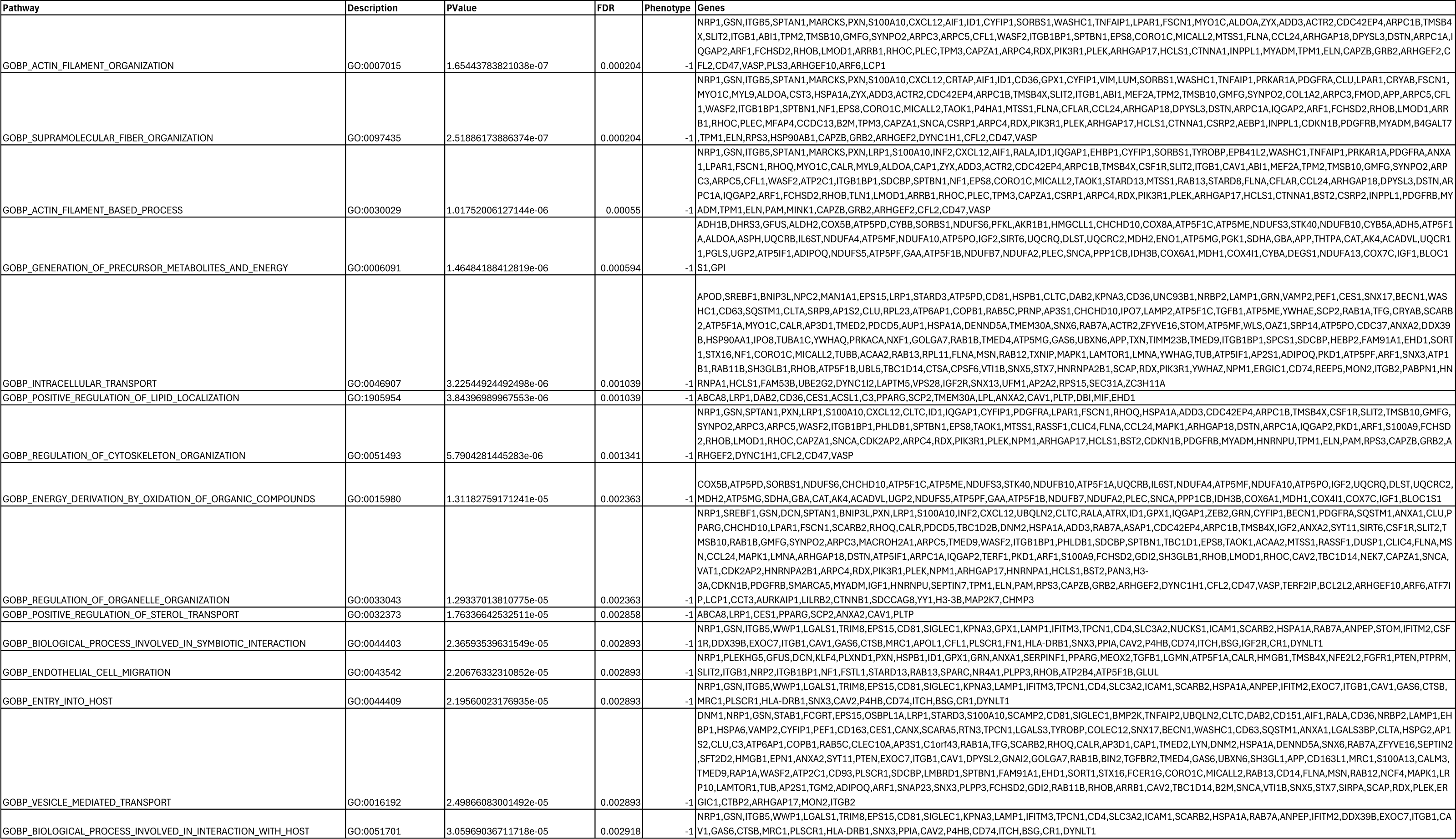
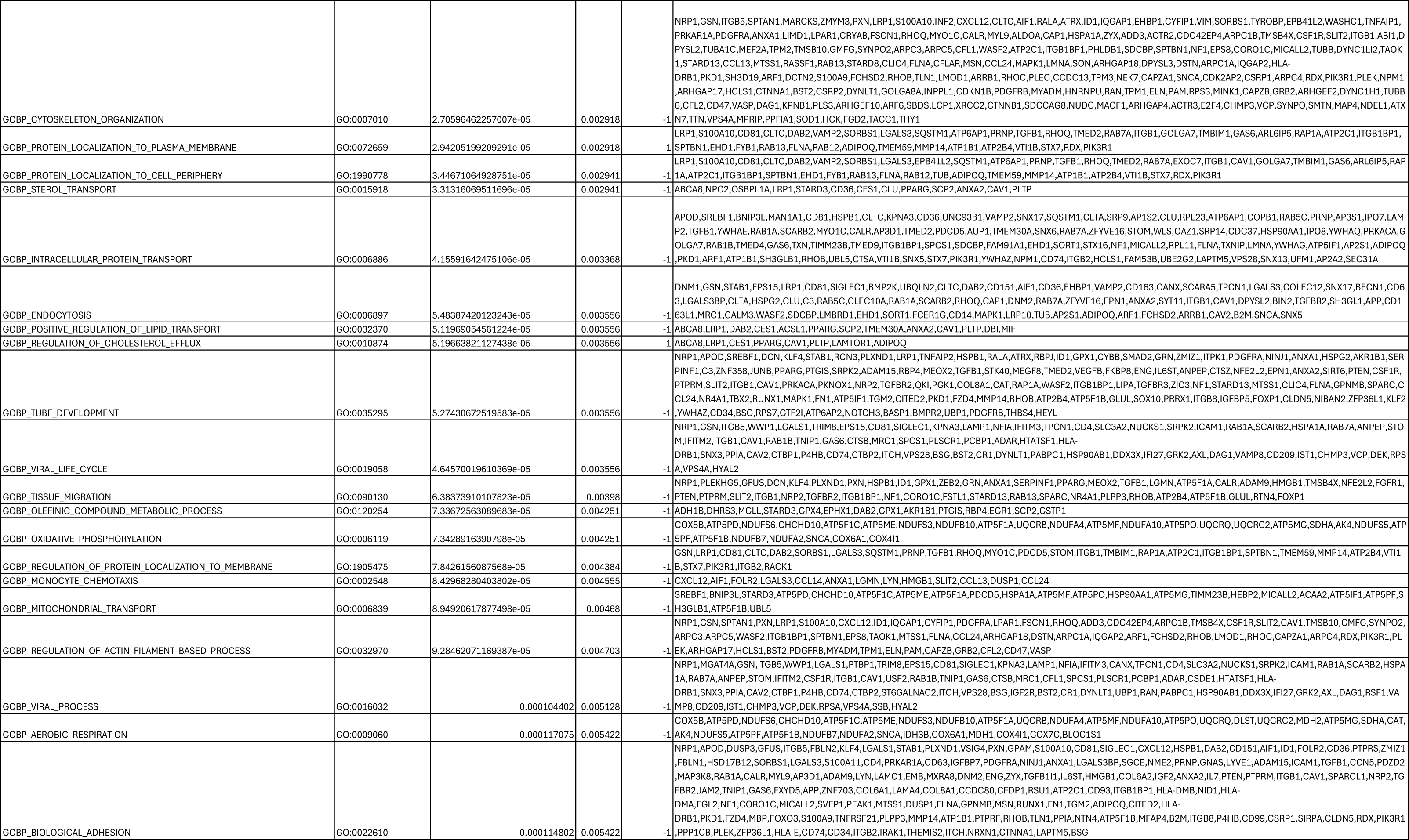
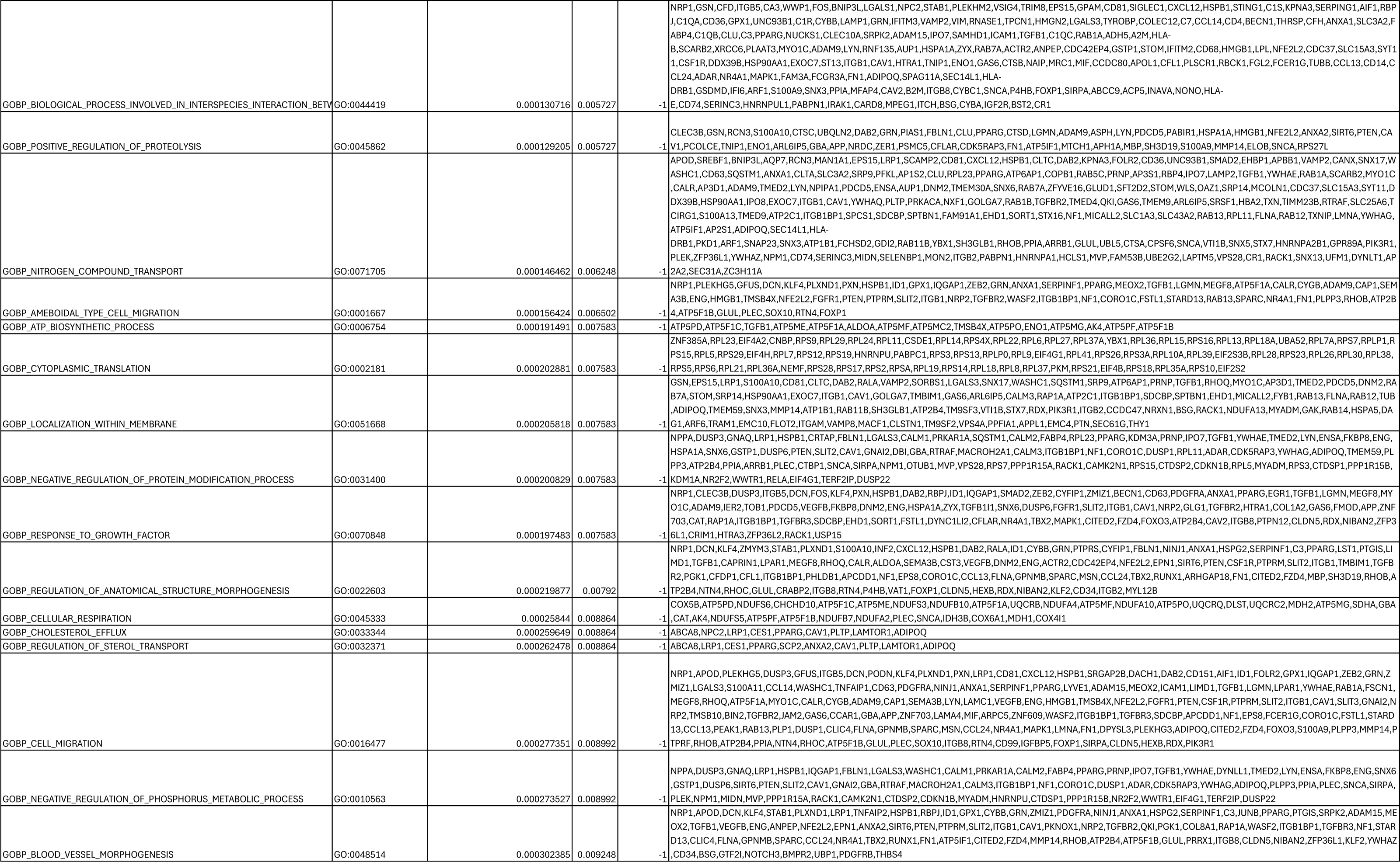
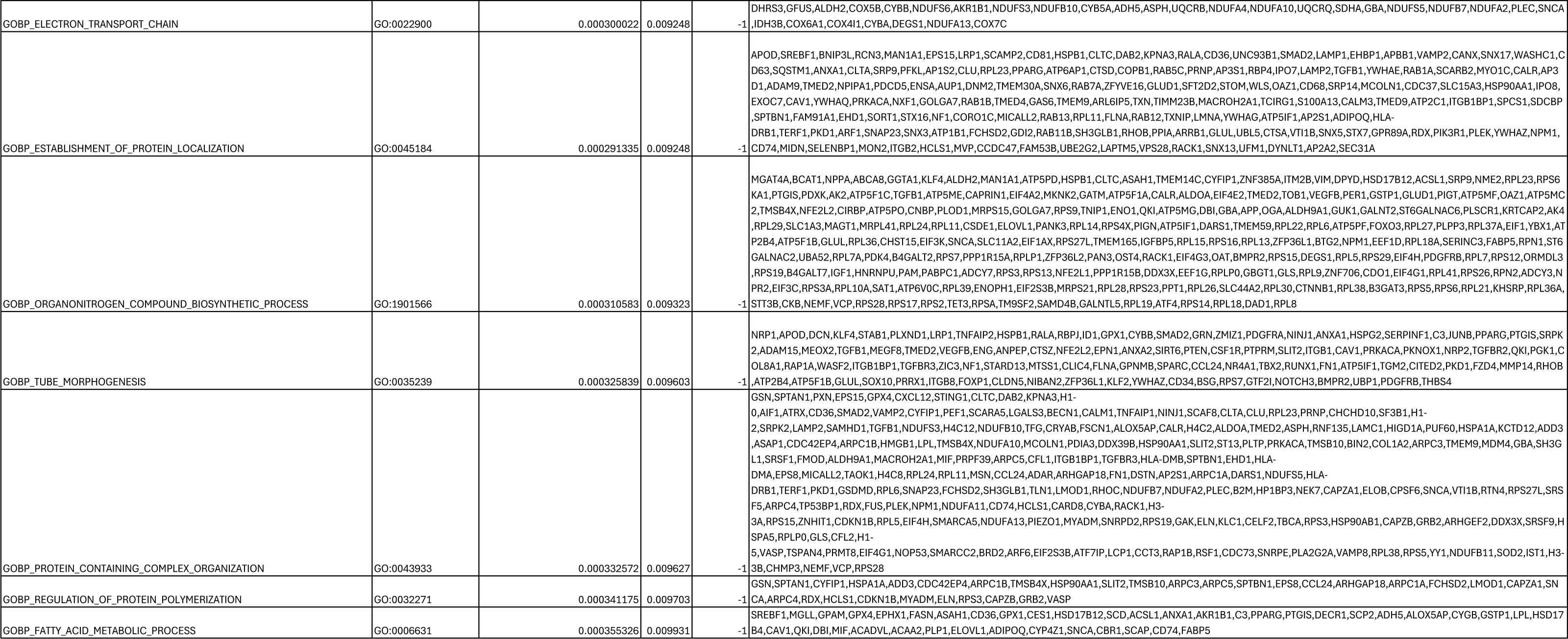

